# Mammalian RAD51 prevents non-conservative alternative end-joining and single strand annealing through non-catalytic mechanisms

**DOI:** 10.1101/768887

**Authors:** Ayeong So, Ali Muhammad, Catherine Chailleux, Laura Sesma Sanz, Sandrine Ragu, Eric Le Cam, Yvan Canitrot, Jean Yves Masson, Pauline Dupaigne, Bernard S. Lopez, Josée Guirouilh-Barbat

## Abstract

The selection of the DNA double-strand breaks (DSBs) repair pathway is decisive for genetic stability/instability. We proposed that it acts according to two successive steps: 1-canonical non-homologous end-joining (C-NHEJ) *versus* single-strand DNA (ssDNA) resection; 2- on ssDNA, gene conversion (GC) *versus* non-conservative single-strand annealing (SSA) or alternative end-joining (A-EJ).

Using intramolecular substrates, we systematically analysed the equilibrium between the different DSB repair pathways. We show that ablation of RAD51 stimulated both SSA and A-EJ but did not stimulate C-NHEJ, validating the two-step model. Moreover, we found that two ATP-mutant dominant-negative forms of RAD51 that stimulated non-conservative repair, failed to load into damaged chromatin, clarifying the role of ATP in RAD51-mediated HR, also. In contrast, another dominant-negative form of RAD51, which retains its DNA binding capacities, repressed SSA and A-EJ, revealing two separable functions of RAD51 i.e. GC and non-conservative repair inhibition. I*n vitro* assays show that the binding of RAD51 on both complementary ssDNA is required to block both spontaneous and RAD52-induced strand annealing. Therefore, RAD51 represses non-conservative repair (SSA and A-EJ), by inhibiting the annealing step through ssDNA occupancy, independently of the catalytic strand-exchange activity required for GC.

## Introduction

DNA double-strand breaks (DSBs) are highly toxic lesions and an important source of genetic instability, which is a hallmark of cancer cells (Negrini *et al*, 2010). On another hand, DSBs are also used to generate beneficial genetic diversity in essential processes such as meiosis, the establishment of the immune repertoire, and neuronal gene expression (for review, see (So *et al*, 2017). Therefore, DSB repair should be tightly controlled to maintain genome stability while allowing for diversity.

Cells use two primary strategies to repair DSBs. The first strategy relies on sequence homology with an intact DNA partner and thus refers to as homologous recombination (HR) including conservative gene conversion (GC). GC is (i) initiated by resection of the DSB producing 3’-single-stranded tails; (ii) the loading of RAD51 onto the ssDNA by BRCA2 generates an ordered ssDNA/RAD51 filament that promotes the central steps of HR: the search for the homologous partner and strand invasion; (iii) DNA synthesis primed on the invading 3’-end; iv) resolution of the intermediate leading to GC with or without crossing over. The second DSB repair strategy, non-homologous end-joining (NHEJ), joins and ligates two DNA double-strand ends without requiring sequence homology; in canonical NHEJ (C-NHEJ), the Ku70/Ku80 heterodimer binds to the DSB and recruits DNA-PKcs, the ligase 4 and its cofactors (Guirouilh-Barbat *et al*, 2004, 2007; Betermier *et al*, 2014; Deriano & Roth, 2013; So *et al*, 2017). In addition to GC and C-NHEJ, both essential for the maintenance of genome stability, other mechanisms exist that exclusively lead to genomic alterations. Single-strand annealing (SSA) is also initiated by resection but anneals the two revealed long complementary ssDNA sequences in a RAD51-independent manner (Haber, 2014; Lambert & Lopez, 2000). In contrast to C-NHEJ, another end-joining (EJ) process, A-EJ (alternative end-joining) also known as alt-NHEJ, B-NHEJ (back-up non-homologous end-joining) or MMEJ (microhomology-mediated end-joining) has been revealed in the absence of KU or XRCC4/Ligase 4 (Guirouilh-Barbat *et al*, 2004, 2007; Rass *et al*, 2009; Betermier *et al*, 2014; Deriano & Roth, 2013; So *et al*, 2017). Like gene conversion and SSA, A-EJ is initiated by ssDNA resection controlled by MRE11/CtIP (Dinkelmann *et al*, 2009; Rass *et al*, 2009; Xie *et al*, 2009); the annealing of complementary micro-homologies (2 to 4 bp), which contrast with the long homologies implicated in SSA, allows the joining of the two resected double-strand ends. Importantly, both SSA and A-EJ are non-conservative processes that inevitably lead to deletions (for review see (Betermier *et al*, 2014; Simsek *et al*, 2011) or translocations (Richardson & Jasin, 2000).

Because HR, SSA and A-EJ are all initiated by resection, we have proposed that the selection of the DSB repair mechanism occurs according to 2 steps (Betermier *et al*, 2014; Rass *et al*, 2009; So *et al*, 2017): 1-competition between C-NHEJ *versus* resection and, 2-on resected DNA ends, competition among HR, A-EJ and SSA. Consistent with this model, defects in C-NHEJ stimulate both gene conversion and A-EJ (Guirouilh-Barbat *et al*, 2007; Delacote *et al*, 2002; Pierce *et al*, 2001; Guirouilh-Barbat *et al*, 2004) and defects in HR stimulate SSA (Stark *et al*, 2004, 2002; Han *et al*, 2017; Lambert & Lopez, 2000). If the regulation of the first step is well documented (including chromatin remodelling and resection initiation or repression), the molecular mechanisms governing the second step, i.e. the selection between GC *versus* SSA or A-EJ, remain less explored and little is known about factors/mechanisms that protect against alternative mutagenic processes SSA and A-EJ.

Here we performed a systematic analysis of different DSB repair pathways (HR, SSA, C-NHEJ and A-EJ), to address the following interconnected questions: 1) What are the consequences of gene conversion invalidation on the equilibrium among other DSB repair processes? 2) What are the molecular bases of the selection of the DSB repair processes at the second step (GC *versus* A-EJ/SSA)? Our data validate the 2-step model and precise molecular bases of the selection at the second step for the choice of the DSB repair pathway in mammalian cells. Indeed, we demonstrated a non-catalytic role of RAD51 for genome stability maintenance: RAD51 prevents both non-conservative SSA and A-EJ by impairing the annealing step of complementary ssDNA. This inhibition acts through RAD51 simultaneous occupancy of both complementary ssDNA strands, and, in living cells, requires the ATPase activity for the loading of RAD51 on damaged DNA, Importantly RAD51-mediated inhibition of SSA and A-EJ does not require the triggering of gene conversion. Collectively, these data reveal an ATP-dependent, but gene conversion-independent role of RAD51 for genome stability maintenance.

## Results

### Silencing RAD51 or BRCA2 stimulates both SSA and end-joining in an epistatic manner

In order to perform a systematic analysis of different DSB repair processes and to validate the two-step model, we used human cells lines containing substrates monitoring gene conversion, SSA or end-joining (EJ) (Figure 1A).

**Figure 1.**
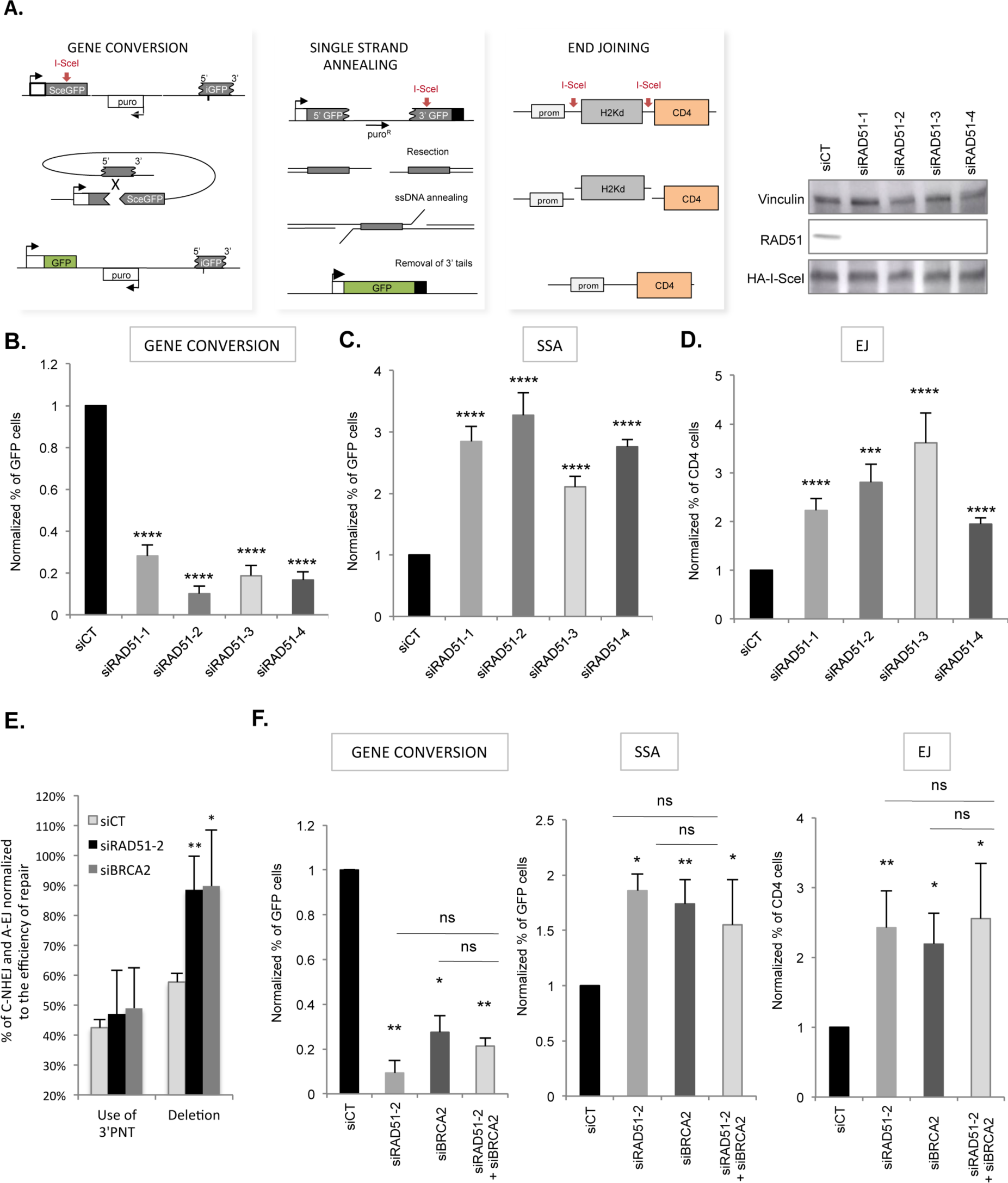
Impact of RAD51 and BRCA2 on the GC *versus* SSA and A-EJ balance. **A.** Three intrachromosomal substrates were used monitoring either gene conversion, SSA or EJ (both C-NHEJ and A-EJ) on a DSB generated by the meganuclease I-SceI. Right panel: Efficiency of the different RAD51 siRNAs and expression of I-SceI. **B, C, and D.** Impact of RAD51 siRNAs on gene conversion (B) SSA (C) and EJ (D). The values are shown normalized to the control siRNA (in black) and represent the average ± SEM of at least 5 independent experiments. **E.** Sequences of the repair junctions from the EJ events after RAD51 or BRCA2 depletion. The use of 3’-PNT (the four 3’-protruding nucleotide generated by I-SceI) mainly corresponds to C-NHEJ, while A-EJ generates deletions at the repair junction (Grabarz *et al*, 2013; Guirouilh-Barbat *et al*, 2004, 2007; Rass *et al*, 2009). **F.** Impact of simultaneous depletion of RAD51 and BRCA2 on gene conversion, SSA, and EJ. The values are shown normalized to the control siRNA (in black) and represent the average ± SEM of at least 5 independent experiments.

First, we verified that silencing RAD51 (4 different siRNAs, Figure 1A) efficiently reduced the levels of GC, as expected (Figure 1B). In parallel, silencing RAD51 also resulted in the stimulation of both SSA and EJ (Figures 1C and 1D). In addition, sequencing of the EJ repair junction revealed an increased frequency of deletions upon RAD51 silencing, but no effect on joining events using the 3’protruding nucleotides (3’PNT) (Figure 1E and Supplemental Figure S1 and S2). Since deletions are a signature of A-EJ and the use of 3’PNT a signature of C-NHEJ (Grabarz *et al*, 2013; Guirouilh-Barbat *et al*, 2004, 2007; Rass *et al*, 2009), these data reveal that RAD51 preferentially suppresses A-EJ rather than C-NHEJ, thus supporting the two-step model for the selection of the DSB repair pathway, rather than a direct competition between C-NHEJ and HR.

Silencing BRCA2 also resulted in the decrease in CG and the stimulation of both SSA and EJ (Figure 1F). Silencing both RAD51 and BRCA2 acted in an epistatic way on GC as well as on SSA and EJ (Figure 1F). Finally, sequencing the EJ junctions, revealed that, similarly to RAD51 silencing BRCA2 stimulated deletions, i.e. mainly A-E-J, rather than C-NHEJ (Figure 1E). As BRCA2 loads RAD51 onto the ssDNA, this suggests that the physical presence of RAD51 onto DNA is required not only to trigger GC, but also to repress both SSA and A-EJ.

Since HR inhibition, through RAD51 ablation, favours SSA and A-EJ, rather than C-NHEJ, this substantiates the two-step DSB repair model. This also supports a role of RAD51 specifically at the second step: in addition to fostering conservative GC, RAD51 also blocks non-conservative A-EJ and SSA, through its loading onto ssDNA.

### Inhibition of GC is not sufficient to stimulate SSA and A-EJ

Because the data above, we addressed whether RAD51 acts on the seletion between GC *versus* SSA and A-EJ, only by channelling DSB repair toward GC and/or, in addition, by physically inhibiting resection and/or annealing through the occupancy of ssDNA ends. Therefore, we used three different dominant-negative forms of RAD51 (DN-RAD51s) that all poison GC (Figure 2A) but with different capacities of binding in damaged chromatin.

**Figure 2.**
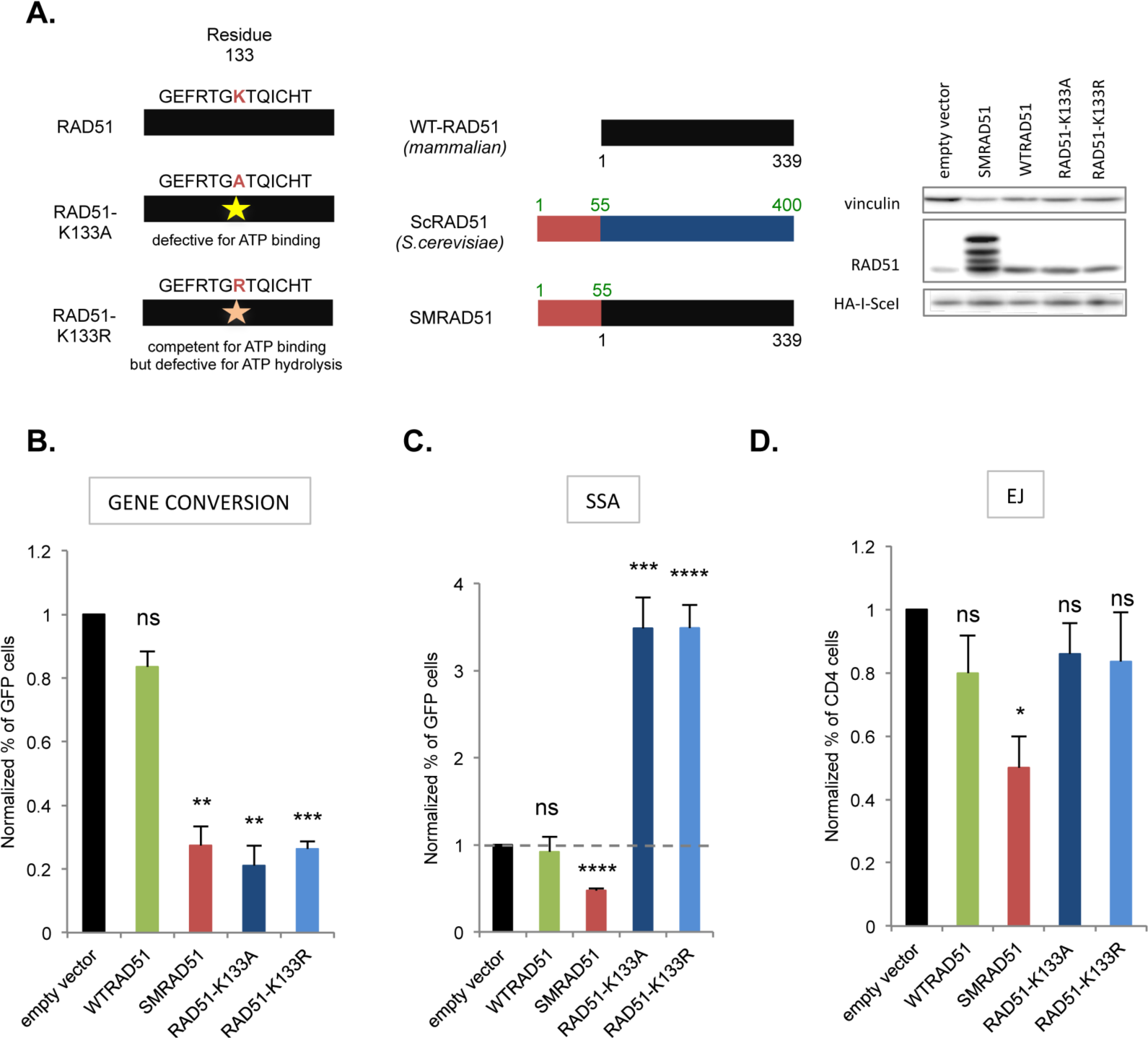
Impact of the different RAD51 dominant negative forms on DSB repair. **A.** Structure of the different RAD51 dominant negative forms. Right panel: The ATPmRAD51s mutated in the ATP binding site. Middle panel: The SMRAD51 chimera. WTRAD51 (mammalian), yeast (*Saccharomyces cerevisiae*) ScRAD51 and the chimera SMRAD51 are aligned, revealing a block of 55 N-ter amino acids (AA) present in the yeast RAD51 but absent from WTRAD51. SMRAD51 corresponds to the fusion of 55 N-ter from yeast to the full-length WTRAD51 (Lambert & Lopez, 2000). Right panel: Expression of the different RAD51 dominant-negative forms. SMRAD51 produces characteristic upper bands (Lambert & Lopez, 2000). **B, C,** and **D.** Impact of the different RAD51 dominant-negative forms on GC **(B),** SSA **(C)** and EJ **(D).** The values are shown normalized to the control transfected with an empty vector (in black) and represent the average ± SEM of at least 5 independent experiments.

Two of the DN-RAD51s are mammalian RAD51 mutated in the ATP binding/hydrolysis site; one mutant (K133A) does not bind ATP and the other (K133R) binds but does not hydrolyse ATP (Stark *et al*, 2002). These ATP-mutant RAD51s (ATPm-RAD51s) are able to bind DNA *in vitro* (Chi *et al*, 2006; Brouwer *et al*, 2018) but are supposed to alter the ssDNA/RAD51 filament activity, through its requirement in the energy from ATP hydrolysis. Another DN-RAD51 corresponds to a yeast and mammalian RAD51 chimera (SMRAD51 from *Saccharomyces*-Mammalian RAD51) (Saintigny *et al*, 2001; Lambert & Lopez, 2001, 2002; Wilhelm *et al*, 2014; Lambert & Lopez, 2000). All 3 DN-RAD51s significantly decreased the efficiency of gene conversion (Figure 2B) in agreement with previous studies (Lambert & Lopez, 2000; Stark *et al*, 2002). Remarkably, although all the DN-RAD51s inhibited gene conversion, we observed different impacts on SSA and EJ (Figure 2C and 2D). Indeed, both ATPm-RAD51s increased SSA (Figure 2C), as reported (Stark *et al*, 2004) and had no impact on the EJ efficiency (Figure 2D). In contrast with RAD51 siRNA and ATPm-RAD51s, SMRAD51, which also poisons GC, decreased SSA and EJ (Figures 2B, 2C and 2D). These data show that inhibition of GC is not sufficient to automatically stimulate SSA and A-EJ. One hypothesis to account for the discrepancies between the different DN-RAD51s proposes that SMRAD51 and the ATPm-RAD51s differ in their capacities to bind damaged DNA in living cells.

### ATP binding and hydrolysis is required for efficient RAD51 loading on damaged DNA, in living cells

We first analysed RAD51 binding on endonuclease-induced DSBs by chromatin-immunoprecipitation (ChIP) (Figure 3A). We used the DIvA system (Iacovoni *et al*, 2010; Caron *et al*, 2012; Aymard *et al*, 2014), in which DSBs are generated upon the nuclear translocation of the restriction endonuclease Asi-SI. Following transfection of an expression vector encoding Flagged-RAD51 (Flag-RAD51), chromatin immunoprecipitation (ChIP) was performed with anti-Flag antibody, and the DNA sequences bound to Flag-RAD51 were quantified by qPCR using specific primers surrounding the cleavage sites. The data showed that SMRAD51 bound to the cleaved DNA sites as efficiently as the wild-type WTRAD51 (Figure 3A). In contrast, RAD51-K133R, poorly bound to the cleaved DNA (Figure 3A).

**Figure 3.**
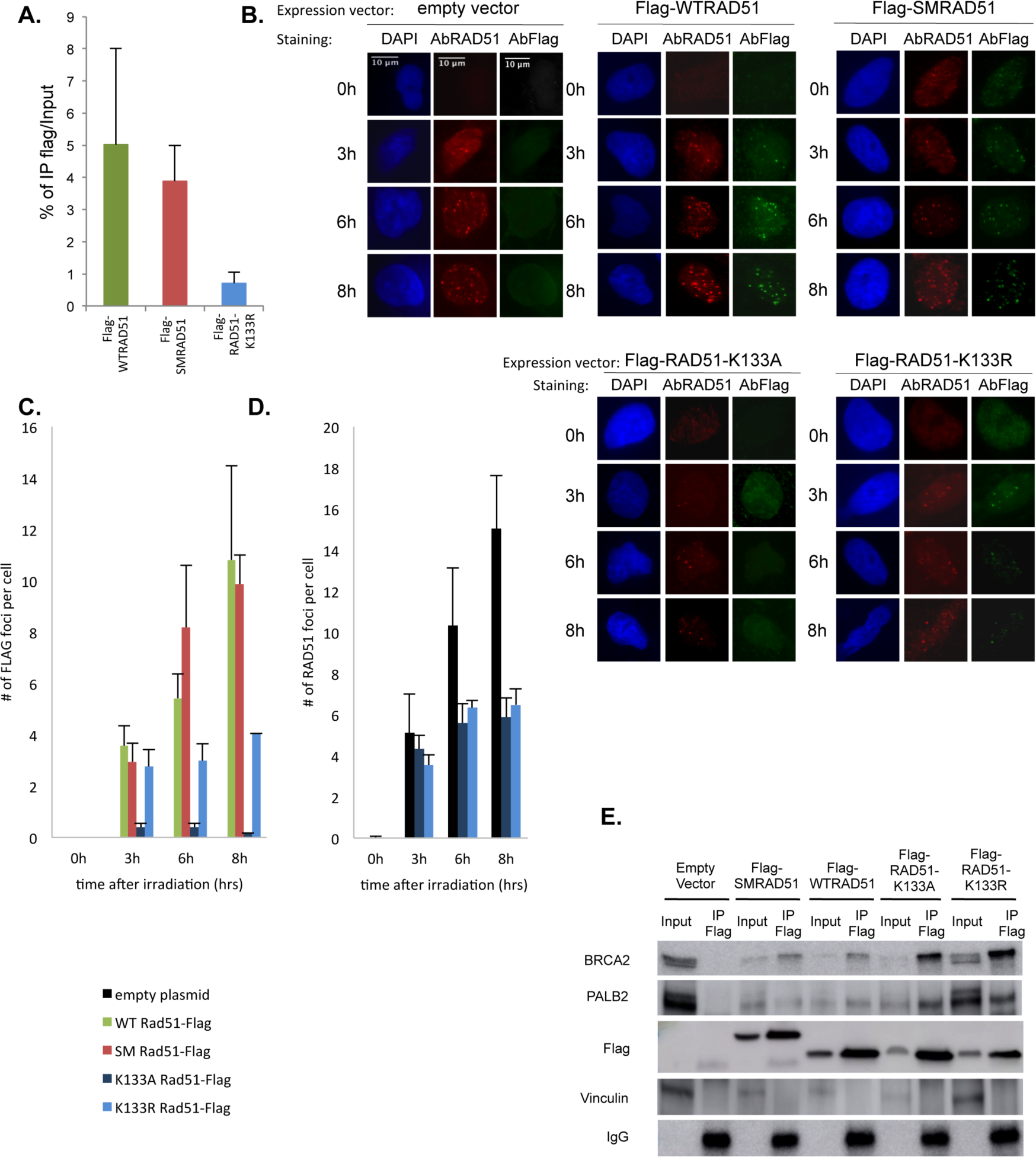
Binding of the different RAD51 dominant-negative forms on damaged DNA. **A.** ChIP for Flagged RAD51s on Asi-SI induced DSBs in DIvA cells. **B.** IR-induced RAD51 foci. Examples of RAD51 foci at different times after IR (6 Gy) revealed either by an anti-Flag antibody. **C.** Kinetics of IR-induced RAD51 foci revealed with an anti-Flag antibody (only exogenous RAD51). The values represent the average ± SEM of at least 5 independent experiments. **D.** Kinetics of IR-induced RAD51 foci revealed with an anti-RAD51 antibody (endogenous + exogenous RAD51). Left panels: Examples of RAD51 foci. Rigth panel: quantification. The values represent the average ± SEM of at least 5 independent experiments. **E.** Co-immunoprecipitation experiments. After transfection with Flag-RAD51s and 4hrs after 6Gy irradiation, cell extracts were immunoprecipitated with a Flag antibody. BRCA2 and PALB2 were then detected by Western blot from the immunoprecipitate using specific antibodies.

We next analysed the formation of RAD51 foci induced by ionizing radiation (IR) (Figure 3B). The anti-RAD51 antibody revealed both endogenous and exogenous RAD51(s). Therefore, we also used Flag-RAD51 to distinguish exogenous RAD51 from the endogenous protein. The two ATPm-RAD51s exhibited defects in IR-induced foci formation (Figures 3B and 3C). Indeed, RAD51-K133A did not assemble into foci and the efficiency of foci formation was strongly decreased for RAD51-K133R (Figure 3C), consistently with the ChIP experiments (Figure 3A). A RAD51 antibody that recognizes both endogenous and exogenous RAD51 revealed that both ATPm-RAD51s affected the efficiency of all RAD51 foci formation, including the endogenous RAD51, accounting for their dominant-negative effect (Figure 3D). In contrast, SMRAD51 efficiently formed IR-induced foci with the same kinetics of assembly/disassembly as endogenous RAD51 or overexpressed WTRAD51 (Figure 3B and 3C).

As the ATPm-RAD51s do not bind efficiently to broken DNA and because BRCA2/PALB2 loads RAD51 onto the ssDNA, we hypothesized that ATP is required for the binding of RAD51 to BRCA2 and PALB2. Then we performed co-immunoprecipitations with the different DN-RAD51s (Figure 3E). All the DN-RAD51s interacted with both BRCA2 and PALB2. These data also show that ATP hydrolysis is not required for the binding of RAD51 to BRCA2 and PALB2. In addition, they suggest that ATPm-RAD51s titrate endogenous proteins such as BRCA2 and PALB2, providing an explanation for the inhibition of endogenous RAD51 foci formation in cells expressing these ATPm-RAD51s. Finally, as ATPm-RAD51 that do not bind DNA stimulate SSA, while SMRAD51 that efficiently binds DNA is capable of inhibiting both SSA and EJ, taken together, these data show that the repression of SSA and A-EJ is correlated to the binding capacities of RAD51 to damaged DNA rather than to the catalysis of GC (compare Figure 2 and 3).

### SMRAD51 specifically disrupts the structure of ssDNA/SMRAD51 filaments

In contrast with ATPm-RAD51s, SMRAD51 efficiently forms foci and binds to cleaved DNA. Nevertheless, SMRAD51 poisons GC. Therefore we hypothesized that the structure of the ssDNA/RAD51 filament, which should be well ordered for efficient HR, might be altered by SMRAD51.

The first, question is whether GC, SSA and A-EJ are poisoned by a mix population of endogenous RAD51 and SMRAD51 or whether SMRAD51 is intrinsically inactive for GC and sufficient to repress A-EJ and SSA. To address this, we used a siRNA against the endogenous RAD51 (targeting the 3’-UTR). SMRAD51 did not rescue GC in RAD51-silenced cells (Figure 4A), suggesting it is intrinsically inactive for HR. Additionally, SMRAD51 retained the capacity to inhibit SSA and EJ in the absence of the endogenous RAD51 (Figures 4B and 4C). Taken together, these data suggest that SMRAD51 binds ssDNA, forming an inactive filament that cannot drive GC but still protects against SSA and A-EJ, giving further evidence of a separation of function between GC and protection against non-conservative DSB repair mechanisms SSA and A-EJ.

**Figure 4.**
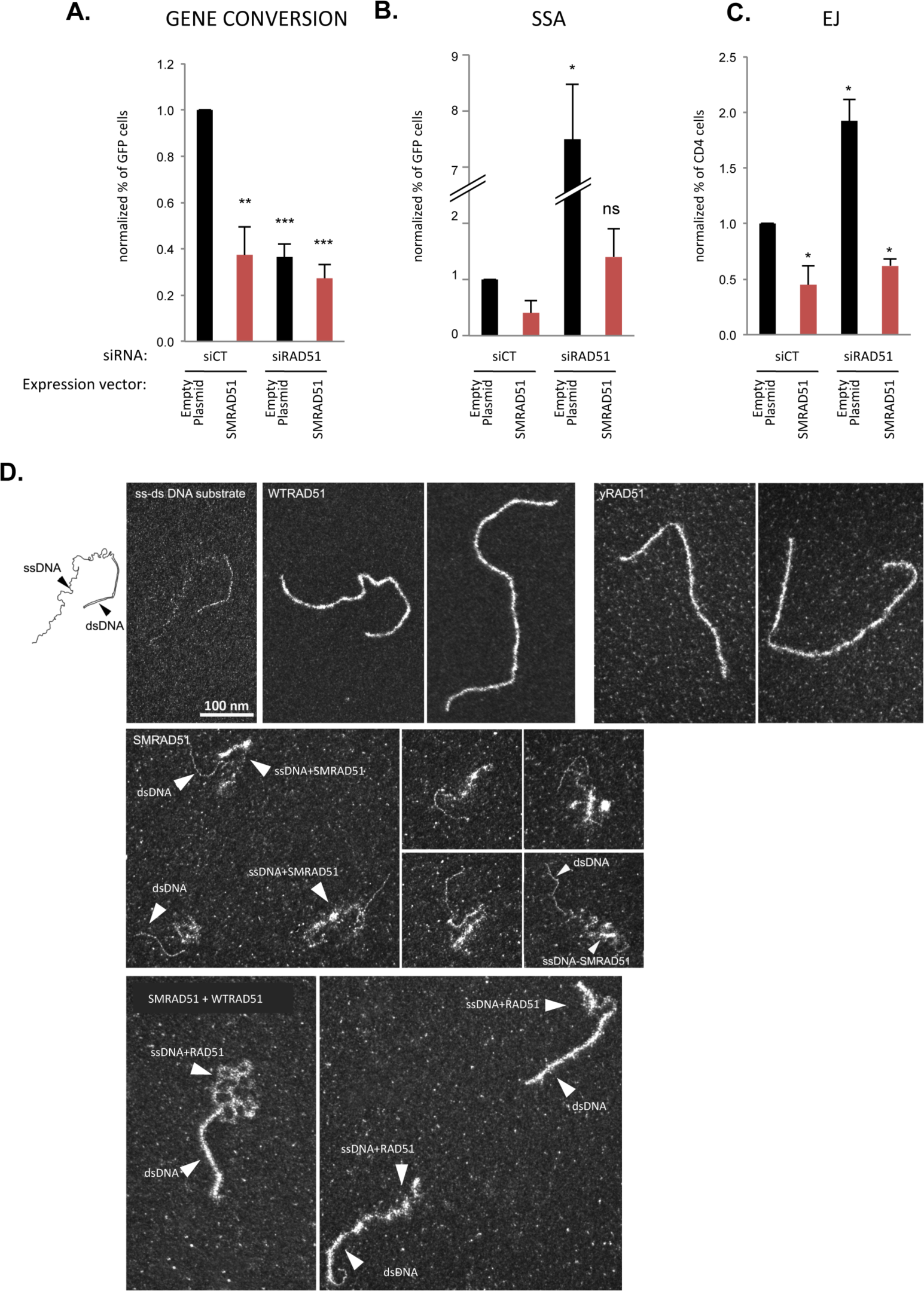
Impact of SMRAD51 on the structure of the ssDNA/RAD51 filament. Impact of mixed endogenous RAD51/SMRAD51 **A)** on gene conversion**, B)** on SSA, **C)** on EJ. SMRAD51 was expressed in cells silencing or not endogenous RAD51. The values are shown normalized to the control siRNA (in black) and represent the average ± SEM of at least 5 independent experiments. **D**. Analysis of the RAD51/ssDNA filament by TEM. Upper-left panels: Substrate used with the single-strand (ss) and double-strand (ds) parts. Upper-middle panel: Filaments formed by the human WTRAD51 protein. Upper-right panels: Filaments form by the *Saccharomyces cerevisiae* yRAD51 protein. Middle panels: Filaments formed by SMRAD51 alone. Lower panels: Filaments formed by a mix of human WTRAD51 and SMRAD51.

To verify these conclusions, we analysed the structure of the ssDNA/RAD51 filament assembled *in vitro* by transmission electron microscopy (TEM). We used a DNA substrate containing one duplex DNA (dsDNA) and one 3’-ssDNA tail (Figure 4D). The human WTRAD51 protein as well as the yeast RAD51 (yRAD51) protein formed ordered filaments. In contrast, the SMRAD51 protein did not bind to dsDNA and formed mis-organized structures on ssDNA. This result is consistent with the data showing that SMRAD51 cannot rescue GC upon endogenous RAD51 silencing but still impairs SSA (see Figures 4A and 4B). Moreover, mixing human WTRAD51 with SMRAD51 resulted in the disruption of the ssDNA/RAD51 filament structure (Figure 4D, lower panel). These data account for the dominant-negative effect of SMRAD51 on GC and underline that it specifically targets the pivotal active species of gene conversion *i.e.* the ssDNA/RAD51 filament.

### RAD51 does not protect against extended resection at endonuclease-induced DSBs

The above data show that in addition to triggering GC, the binding on DNA of RAD51 (even inactive for HR) also impairs non-conservative DSB repair processes, which involve both resection and annealing of complementary ssDNA. Therefore, this suggests that RAD51 either protects against resection, as shown for arrested replication forks (Schlacher *et al*, 2011; Ying *et al*, 2012), and/or impairs the annealing of complementary ssDNA, through ssDNA occupancy. We addressed the resection hypothesis using the DIvA system that allows to quantify the resection at different distances of Asi-SI cutting sites in the genome (Zhou *et al*, 2013; Cohen *et al*, 2018). We measured resection on two different chromosomes (chromosomes 1 and 20) that have been previously mapped (Zhou *et al*, 2013; Cohen *et al*, 2018). As expected resection decreased with the distance from the Asi-S1-cleaved sites. Moreover, as a control, silencing CtIP, which is involved in resection initiation, significantly affected the resection efficiency (Figure 5A). Remarkably, silencing RAD51 did not significantly impact the efficiency of resection at either site (Figure 5A). This result is consistent with the fact that silencing RAD51 did not affect the size distribution of deletions at the A-EJ repair junction (Supplementary data S3).

**Figure 5.**
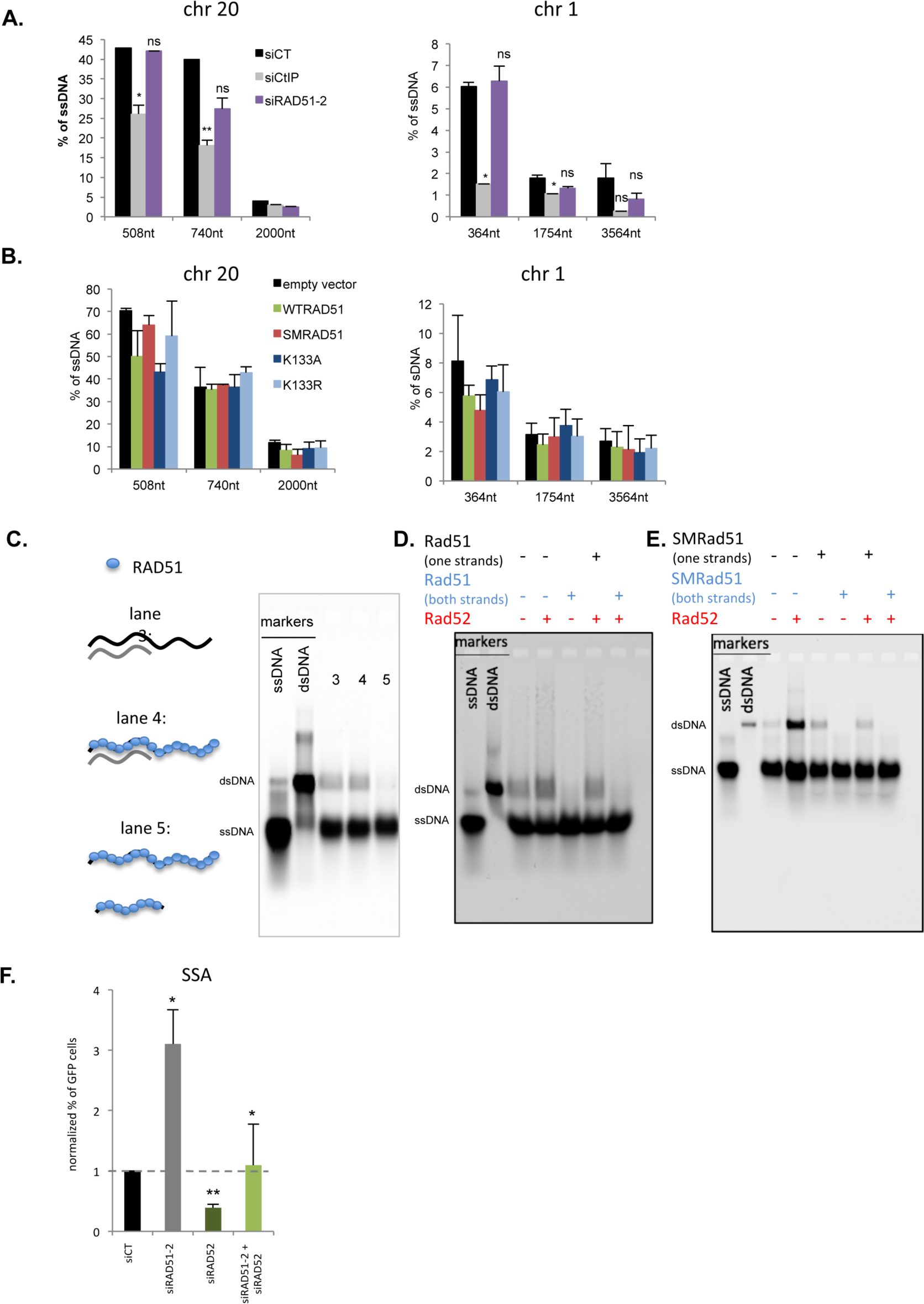
Impact of RAD51 on resection and annealing. **A.** Impact of silencing RAD51 on resection. Two Asi-SI sites, on chromosome 20 and 1 were analyzed, using either BamH1 (chromosome 1) or BanI (Chromosome 20) endonucleases. The values are the average ± SEM of at least **5** independent experiments. **B.** Impact of the dominant-negative forms of RAD51 on resection. Two AsiSI sites (chromosome 20 and 1) were analyzed. The values are the average ± SEM of at least **5** independent experiments. **C.** Impact of RAD51 on the annealing of two complementary ssDNAs. Lanes 1 and 2 corresponds to markers (ssDNA or dsDNA). Lane 3: Incubation of two naked complementary ssDNAs. Spontaneous annealing gives a band migrating at the position of the dsDNA. Lane 4: One ssDNA was coated with RAD51 prior incubation with its complementary partner. Lane 5: Both complementary ssDNAs were coated with RAD51 prior their co-incubation. **D.** Impact of RAD51 on the annealing of two complementary ssDNAs stimulated by RAD52. **E**. Impact of SMRAD51 on the annealing of two complementary ssDNAs stimulated by RAD52. **F.** Impact of RAD52 *versus* RAD51 on SSA. The values are shown normalized to the control siRNA (in black) and represent the average ± SEM of at least 3 independent experiments.

Moreover, none of the dominant-negative forms of RAD51 affected the resection efficiency (Figure 5B). Taken together, these data suggest that RAD51 does not protect against long resections (hundreds of bp) at DSBs generated by endonucleases.

### RAD51 prevents RAD52-mediated annealing of complementary ssDNAs, when loaded on both complementary strands

We addressed the potential impact of RAD51 on annealing using an *in vitro* approach with purified proteins (Figure. 5C-E). *In vitro*, two complementary ssDNAs spontaneously anneal, forming a DNA duplex (dsDNA). Loading RAD51 onto one ssDNA strand did not impact the spontaneous annealing efficiency, while loading RAD51 on both complementary ssDNAs abolished it (Figure 5C). As expected, adding RAD52 stimulated the annealing of 2 complementary naked ssDNA (Figure 5D and 5E). RAD51 loaded on both complementary ssDNA also inhibited RAD52-mediated annealing (Figure 5D). Moreover, SMRAD51 completely abolished both spontaneous and RAD52-mediated annealing, when loaded on both ssDNA strands, and decreased it when loaded on only one strand (Figure 5E).

Collectively, these data show that RAD51 should occupy DNA simultaneously on both complementary strands to prevent both spontaneous and RAD52-mediated annealing, in a process that does not require to triggering strand exchange.

As SSA require the annealing of long homologies, the above data suggest that the stimulation of SSA observed upon RAD51 depletion (see Figure 1 C-F and 2C-D) might rely on RAD52. Consistent with the role of RAD52 in SSA, RAD52 silencing did decrease SSA efficiency (Figure 5F). Additionally, silencing RAD52 abolished the stimulation of SSA resulting from RAD51 silencing (Figure 5F).

## Discussion

Our data bring molecular support to the two-steps model for the selection of the DSB repair pathway (Betermier *et al*, 2014; Rass *et al*, 2009; So *et al*, 2017). Particularly, we show here that RAD51 plays a pivotal role at the second step, through at least two separable mechanisms: i) channelling DSB repair toward HR, and ii) GC-independent inhibition of the annealing of complementary sequences. The increase of A-EJ and SSA upon HR deficiency has been separately reported in different studies (Stark *et al*, 2004; Ahrabi *et al*, 2016; Han *et al*, 2017), always proposing that the DSB repair was redirected to alternative mechanisms due to the lack of the RAD51 strand exchange activity. Here, we show that inhibiting GC with the dominant-negative SMRAD51 does not stimulate SSA and EJ. Thus, the stimulation of SSA and A-EJ in the absence of RAD51 is not a direct consequence of the inhibition of GC itself, as previously proposed, but rather results from the absence of repression of the annealing of complementary ssDNA by RAD51. In budding Yeast, Rad51 suppresses Rad52-dependent ssDNA annealing, facilitating DNA strand invasion *in vitro* (Wu *et al*, 2008; Sugiyama & Kantake, 2009). Here we show that GC and repression of annealing are separable processes: indeed SMRAD51 that is inactive for GC but binds DNA, retains the full capacity to inhibit annealing, *in fine* repressing both SSA and A-EJ. More specifically, we show here that the binding of RAD51 on both single stranded ends is required to prevent the annealing. This characteristic should be important to preserve the capture of the second end during GC, as discuss below.

*In vitro,* ATPm-RAD51 proteins bind DNA (Chi *et al*, 2006; Brouwer *et al*, 2018). Therefore, it was proposed that they exert their dominant-negative effects by preventing the ssDNA/RAD51 filament from achieving its energy needs. We show here that in living cells, ATPm-RAD51s are inefficiently loaded onto damaged DNA, suggesting that the primary role of ATP is in fact the transfer of RAD51 to the damaged DNA.

We also show here that the main SSA/A-EJ step blocked by RAD51 is the annealing step, but not the long resections at least necessary for SSA. In contrast, RAD51 has been shown to protect arrested replication forks from resection (Schlacher *et al*, 2011; Ying *et al*, 2012). This suggests that the protection of arrested replication forks and of endonuclease-induced DSBs is differently regulated.

PolQ has been shown to stimulate A-EJ and to remove RAD51 from the DNA (Ceccaldi *et al*, 2015). This is highly consistent with our data and with the central role of RAD51 in the second-step choice in DSB repair.

The present data are summarized in the model in Figure 6. At the first step C-NHEJ competes with resection; at the second step, BRCA2/PALB2 load RAD51 onto the resected ssDNA in an ATP-dependent manner. Then, RAD51 (i) triggers GC and (ii) blocks the annealing of complementary ssDNA strand, inhibiting both SSA and A-EJ. Note that the choice between SSA and A-EJ should be influenced by the presence or not of long complementary ssDNA, which are much less frequent than micro-homologies. During RAD51-dependent GC process, RAD52 can also act on the annealing step, allowing the capture of the displaced strand by the resected second double-strand ends (Miyazaki *et al*, 2004; McIlwraith & West, 2008; Sugiyama *et al*, 2006; Brouwer *et al*, 2017). We show here that RAD51 blocks annealing when it is loaded on both complementary ssDNAs, which is the situation in both SSA and A-EJ. In contrast, RAD51 does not efficiently block annealing when it is loaded on only one ssDNA, which occurs for the capture of the second DSB during gene conversion. Therefore this specificity of RAD51 allows the protection against non-conservative SSA and A-EJ, while allowing conservative GC.

**Figure 6.**
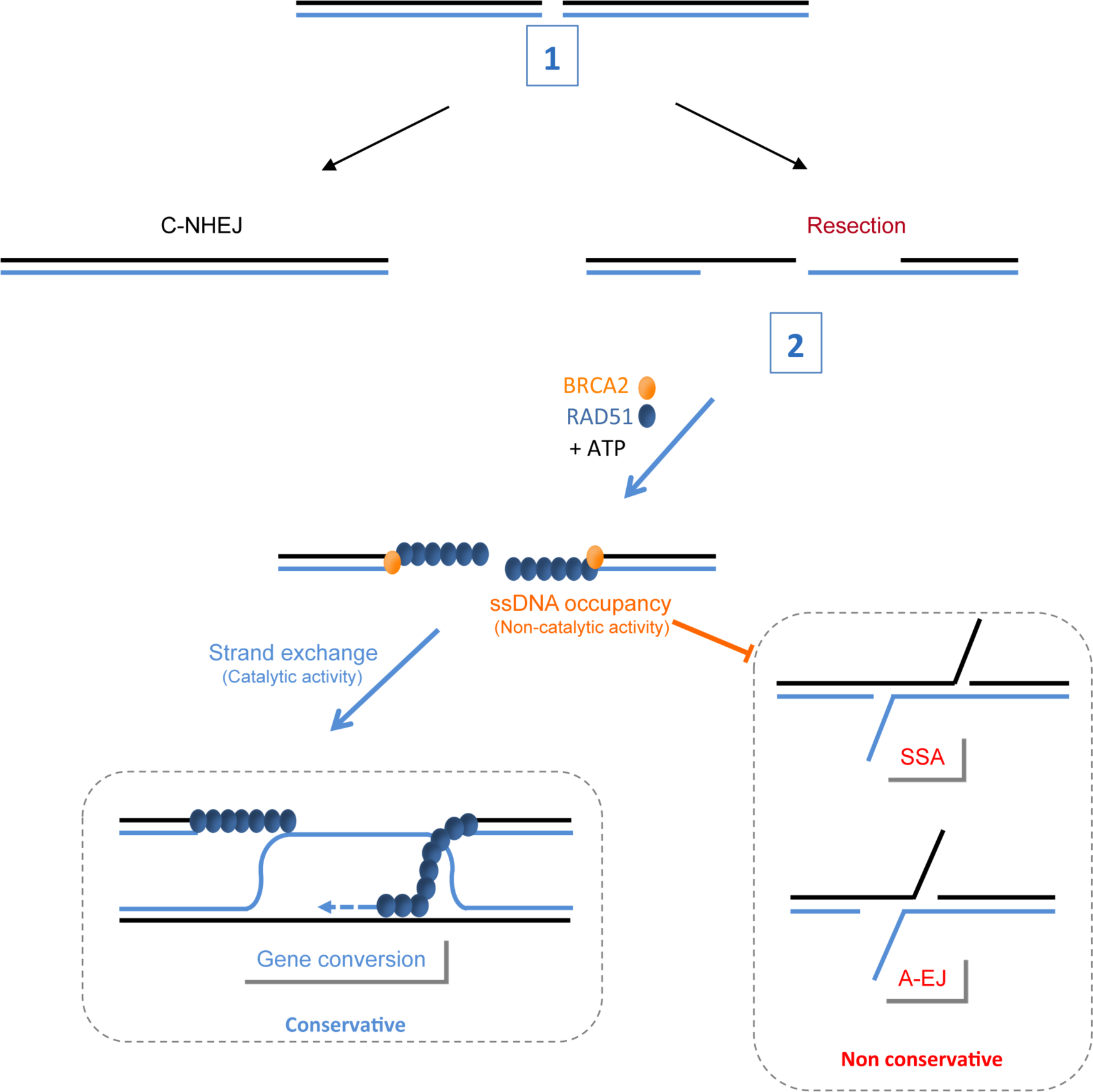
The roles of RAD51 for the selection of the DSB repair pathway. **1.** C-NHEJ favoured by 53BP1 and Ku86/70 competes with resection on DSBs. **2.** After resection, BRCA2 loads RAD51 onto the ssDNA in an ATP-dependent manner. The occupancy of the ssDNA by RAD51 triggers gene conversion (in a catalytic manner) and suppresses the annealing step (in a non-catalytic manner), which could be RAD52-dependent, of non-conservative SSA and A-EJ. The blockage of annealing by RAD51 does not require the strand exchange activity. Note that RAD52 might cooperate with RAD51 during GC in the capture of the second double-strand end (left panel).

Altogether our data detail the switch between the cooperation and antagonism of RAD51 and RAD52 for the maintenance of genome stability in mammalian cells: RAD51 loaded on both single-stranded DNA prevents RAD52 mediated mutagenic DSB repair process(es), however RAD51 and RAD52 also cooperate for gene conversion when RAD51 is loaded on only one single-stranded DNA.

## Materials and Methods

### Cells

We used cell lines with intrachromosomal substrates monitoring either gene conversion, SSA, EJ (both C-NHEJ and A-EJ) after targeted induction DNA double strand breaks by the meganuclease I-SceI were used. The RG37 cell line (Dumay *et al*, 2006) was derived from SV40-transformed GM639 human fibroblasts and stably contain the pDR-GFP, a gene conversion reporter, which restores a functional GFP gene upon I-SceI cleavage (Pierce *et al*, 1999). GC92 cells (Rass *et al*, 2009) are also derived from SV40-transformed GM639 human fibroblasts and contain the pCOH-CD4/CD4-3200bp (je pense qu’on doit choisir un nom et pas deux : donc soit on recombine “pCOH-CD4-3200bp” soit on choisit un des deux) substrate that monitors EJ by the expression of the membrane antigen CD4. Because I-SceI cleaves in two non-coding sequences, both error-prone and error-free repair are measured; *i.e.*, both C-NHEJ (conservative repair) and A-EJ (exclusively mutagenic repair) are recorded. Sequence of the repair junction allows the estimation of the C-NHEJ/A-EJ ratio (Grabarz *et al*, 2013; Guirouilh-Barbat *et al*, 2004, 2007; Rass *et al*, 2009). U2OS-SSA cells were previously described (Gunn *et al*, 2011; Gunn & Stark, 2012). The DIvA cell line (Asi-SI-ER-U2OS) is a U2OS cell line (human osteosarcoma) previously established and described by Iacovoni et al (2010). DNA double strand breaks are induced at located regions in the genome by the Asi-SI endonuclease. Asi-SI is sequestered in the cytoplasm, and after addition of hydroxy tamoxifen (300 nM) to the culture medium for 4 hours, Asi-SI translocates into the nucleus and cuts DNA. All cells were cultured in DMEM supplemented with 10% foetal calf serum (FCS) and 2 mM glutamine and were incubated at 37°C with 5% CO_2_.

### Transfection

The meganuclease I-SceI was expressed by transient transfection of the pCMV-HA-I-SceI expression plasmid (Liang *et al*, 1998) with Jet-PEI according to the manufacturer’s instructions (Polyplus transfection). The expression of HA-tagged I-SceI was verified by Western blotting, in each condition. For silencing experiments, 50,000 cells were seeded 1 day before transfection; these experiments were performed using INTERFERin following the manufacturer’s instructions (Polyplus Transfection) with 20 nM of one of the following siRNAs: Control (5’-AUGAACGUGAAUUGCUCAA-3’), RAD51-1 (cat# L003530-00-0010, Dharmacon), RAD51-2 (5’-GAAGCUAUGUUCGCCAUUA-3’), RAD51-3 (3’-UTR, 5’GACUGCCAGGAUAAAGCUU-3’), RAD51-4 (3’UTR, 5’-GUGCUGCAGCCUAAUGAGA-3’), BRCA2 (5’-GCUGAUCUUCAUGUCAUAA-3’) or RAD52 (5’-CCAACGCACAACAGGAAAC-3’), all of which (except those ordered from Dharmacon) were synthesized by Eurofins. Forty-eight hours later, the cells were transfected with the pCMV-HA-I-SceI expression plasmid.

### Measure of gene conversion, SSA and EJ efficiency by FACS

After transfection with the pCMV-HA-I-SceI plasmid and incubation for 72 hours, the cells were collected 50 mM EDTA diluted in PBS, pelleted and fixed with 2% paraformaldehyde for 20 minutes. The percentage of GFP-expressing cells was scored by FACS analysis using a BD Accuri C6 flow cytometer (BD). The percentage of CD4-expressing cells was measured after incubation for 10 minutes with 1 µl of anti-CD4 antibody coupled to Alexa 647 (rat isotype, RM4-5, Invitrogen). For each cell line, at least 3 independent experiments were performed, and HA-I-SceI expression and silencing efficiency were verified each time by Western blot.

### Western blotting

For Western blot analysis, the cells were lysed in buffer containing 20 mM Tris HCl (pH 7.5), 1 mM Na_2_EDTA, 1 mM EGTA, 150 mM NaCl, 1% (w/v) NP40, 1% sodium deoxycholate, 2.5 sodium pyrophosphate, 1 mM b-glycerophosphate, 1 mM NA_3_VO_4_ and 1 µg/ml leupeptin supplemented with complete mini protease inhibitor (Roche). Denatured proteins (20-40 µg) were electrophoresed in 9% SDS-PAGE gels or MiniPROTEAN® TGX™ 4-15% Precast gels (BIORAD), transferred onto a nitrocellulose membrane and probed with specific antibodies, including anti-Vinculin (1/8,000, SPM227, Abcam), anti-RAD51 (1/1,000, Ab-1 Millipore), and anti-HA (1/1,000, HA.11 clone 16B12, Covance). Immunoreactivity was visualized using an enhanced chemiluminescence detection kit (ECL, Pierce).

### Junction sequence analysis

We amplified the junction sequences through PCR of genomic DNA using the CMV-6 (5’-TGGTGATGCGGTTTTGGC-3’) and CD4-int (5’-GCTGCCCCAGAATCTTCCTCT-3’) primers. The predicted size of the PCR product is 732 nt. The PCR products were cloned with a TOPO PCR cloning kit (Invitrogen Life Technologies) and sequenced (Eurofins). For each sample, 2 to 5 independent experiments were analyzed. In each of these experiments, HA-I-SceI expression and the silencing efficiency were verified by Western blot.

### Immunofluorescence

Cells were seeded onto slides and transfected with empty vector, Flag-SMRAD51, Flag-WTRAD51, Flag-RAD51 K133A and Flag-RAD51 K133R. Seventy-two hours after transfection, the cells were washed with PBS, treated with CSK buffer (100 mM NaCl, 300 mM sucrose, 3 mM MgCl_2_, 10 mM Pipes pH 6.8, 1 mM EGTA, 0.2X Triton, and protease inhibitor cocktail (cOmplete ULTRA Tablets, Roche)) and fixed in 2% paraformaldehyde for 15 min. The cells were then permeabilized in 0.5% Triton-X 100 for 10 min, saturated with 2% BSA and 0.05% Tween20 and probed with anti-Flag (1/400, F3165 Sigma-Aldrich) or anti-RAD51 (1/500, PC130, Merck Millipore) antibody for 2 h at RT or overnight at 4°C. After 3 washes in PBS-Tween20 (0.05%) at RT, the cells were probed with Alexa-coupled anti-mouse or anti-rabbit secondary antibody (1/1,000, Invitrogen) for 1 h at RT. After 3 washes, the cells were mounted in DAKO mounting medium containing 300 µM DAPI and visualized using a using a fluorescence microscope (Zeiss Axio Observer Z1) equipped with an ORCA-ER camera (Hamamatsu). Image processing and foci counting were performed using the ImageJ software.

### Co-immunoprecipitation

GC92 cells grown on 75 cm^2^ cell culture flasks were irradiated with an irradiator (XRAD320, 6 Gy) for 4 h before collection. Cells were washed in PBS and pellets were resuspended with 300 µl lysis buffer (150 mM NaCl, 25 mM Tris pH 7.5, 1 mM EDTA, 0.5% NP40, protease inhibitors (cOmplete ULTRA Tablets, Roche) and phosphatase inhibitors (phosphatase inhibitor cocktail 2/3, P5726/P0044, Sigma)) and then incubated for 1 h at 4°C. Cell extracts were centrifuged at 13,200 rpm for 20 min at 4°C. Following the measurement of the total protein amount, the supernatant fraction was incubated with 15 µl fetal bovine serum for 1 h at 4°C and then incubated with 10 µl pre-cleaned magnetic beads at 12 rpm for 30 min at 4°C. Next, 200 µg protein were incubated with DNase I (7,5 U) and 1 µg anti-Flag resin (cat# F3165, Sigma-Aldrich) at 12 rpm overnight at 4°C. Following an incubation with pre-cleaned magnetic beads at 12 rpm for 4 h at 4°C, the resin was washed three times with lysis buffer. Laemmli 2X (62.5 mM Tris-HCl, pH 6.8, 10% glycerol, 1% LDS, and 0.005% bromophenol blue) was added and the proteins were boiled for 5 min at 95°C. Denatured protein extracts were resolved using 4-9% SDS-PAGE (4–15% Mini-PROTEAN® TGX™ Precast Protein Gelscat# 4561083, Biorad) and then transferred to a nitrocellulose membrane.

### ChIP

DIvA cells (25 × 10^6^) were transfected by electroporation (Amaxa solution V) using 20 µg of plasmid coding for the different forms of RAD51. Cells were seeded in a plate; 24 h later, the cells were treated or not with hydroxy tamoxifen (300 nM final concentration) for 4 h. The cells were crosslinked with formaldehyde (1%, 20 min) followed by cell lysis, DNA sonication and immunoprecipitation of the protein/DNA complexes with specific antibodies (anti-Flag M2, Sigma) or without antibodies as a negative control. DNA/protein complexes were collected with a mix of protein G and A agarose beads. Crosslinking was reversed by the addition of NaCl and then samples were treated with RNase A and proteinase K and DNA purified by centrifugation using a GFX PCR column (Amersham). DNA was quantified by qPCR using the SYBR Green qPCR master mix (Biorad) and the following specific primers: (1) DSB site, 5’-GATTGGCTATGGGTGTGGAC-3’and 5’-CATCCTTGCAAACCAGTCCT-3’ and (2) control, 5’-GGCGACCTGGAAGTCCAACT -3’ and 5’-CCATCAGCACCACAGCCTTC -3’.

### TEM experiments

For TEM analysis, RAD51 filaments were examined by incubating 15 µM of 5’-DNA junction substrate (400 bp double-stranded DNA with a 1000 nt single-stranded DNA overhang) with 5 µM RAD51 (WTRAD51, ScRAD51, SMRAD51 or both) for 3 min at 37°C. Then, 0.1 µM RPA (WTRPA or ScRPA) was added for 10 min at 37°C. The buffer for the reactions with WTRAD51 and SMRAD51 was 10 mM Tris-HCl (pH 8), 50 mM sodium chloride, 2 mM calcium chloride, 1 mM DTT and 1 mM ATPγ, whereas the buffer for the ScRAD51 filament reaction was 10 mM Tris-HCl (pH 8), 50 mM sodium chloride, 3 mM magnesium chloride, 1 mM DTT and 1.5 mM ATP. The reactions were quickly diluted 100× in a buffer containing 10 mM Tris-HCl pH 8, 5 mM MgCl_2_, and 50 mM NaCl. For one minute, a 5 μL drop was deposited on a 600-mesh copper grid previously covered with a thin carbon film and activated with pentylamine by glow-discharge using the Dubochet device. Grids were rinsed and positively stained with aqueous 2% (w/v) uranyl acetate, dried carefully with a filter paper and observed in annular dark-field mode using a Zeiss 902 transmission electron microscope. Images were captured at a magnification of 85,000× with a MegaviewIII CCD camera and analyzed with the iTEM software (Olympus Soft Imaging Solution).

### Resection assay

Resection measurements on DIvA Cells were performed as previously described (Zhou *et al*, 2013). Briefly, after hydroxy tamoxifen treatment cells were collected for genomic DNA extraction (DNeasy blood & tissue kit, Qiagen), 100-200 ng genomic DNA was treated with 16 U of the appropriate restriction enzyme overnight at 37°C. After enzyme inactivation, the digested genomic DNA was used for qPCR (TAKARA mix for qPCR) with the primers listed in the table below.

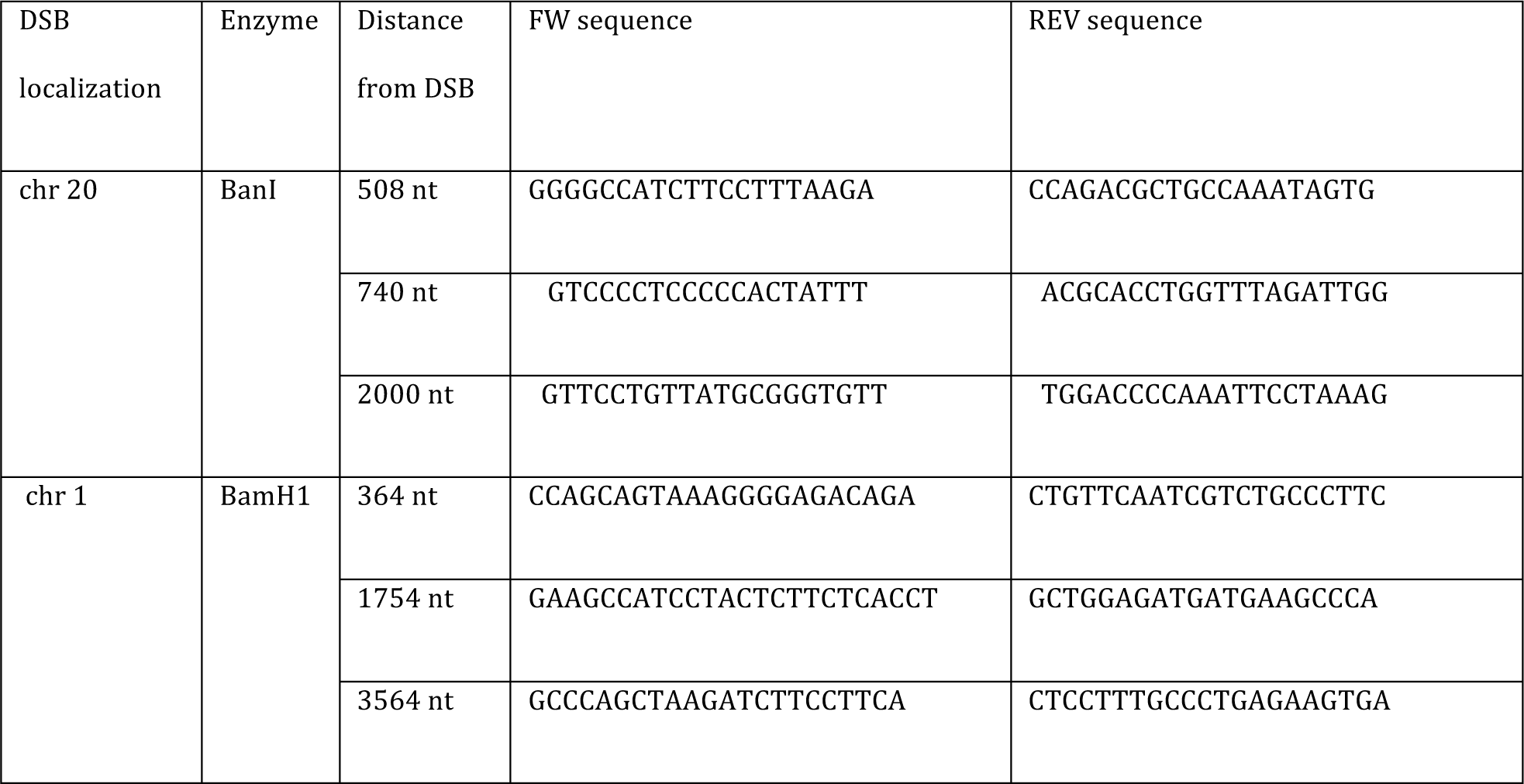

The percentage of ssDNA was calculated with the following equation: ssDNA % = 1/(2^(△Ct-1) + 0.5)*100, where △Ct = Ct of the digested sample – Ct of the non-digested sample.

### Single strand annealing assay

All reactions were performed in a buffer containing 10 mM Tris-HCl (pH 8), 50 mM sodium chloride, 2 mM calcium chloride, 1 mM DTT and 1 mM ATPγs. In reaction A, 0.1 µM (8 µM nucleotides) of a 178-mer oligonucleotide (5’-ATCATCACCATCACCATTGAGTTTAAACCCGCTGATCAGCCTCGACTGTGCCTTCTAGTTGCCAGCCATCTGTTGTTTGCGGTTCCCAACGATCAAGGCGAGTTACATGATCCCCCATGTTGTGCAAAAAAGCGGTTAGCTCCTTCGGTCCTCCGATCGTTGTCAGAAGTAAGTTGGC-3’) was pre-incubated with 2.6 or 8 µM WTRAD51 (ratio protein/nt: 1/3 or 1/1, respectively) for 10 min at 37°C in a final volume of 5 μl. In reaction B, 0.1 µM of a 3′-Cy5-labeled 80-mer oligonucleotide, which is complementary to the 5′-portion of the 178-mer described in reaction A (5’-GCAAACAACAGATGGCTGGCAACTAGAAGGCACAGTCGAGGCTGATCAGCGGGTTTAAACTCAATGGTGATGGTGATGAT-Cy5-3’) was pre-incubated under the same conditions with 2.6 or 8 µM WTRAD51 (ratio protein/nt: 1/3 or 1/1, respectively). Reactions A and B were mixed together for 5 min at 30°C. Then, 5 µM of an unlabeled DNA (80-mer) was added to the reaction. The total reaction was stopped by the addition of 1% SDS (w/v) and 25 mM EDTA and deproteinized (30 min incubation at 37°C with 2 mg/ml proteinase K). Samples were run in a 3% (w/v) agarose gel at 80 V for 30 min in 0,5x TAE buffer. Fluorolabeled DNA species were visualized by fluorescent imaging using a Typhoon FLA 9500 (GE Healthcare Life Sciences).

### Statistical Analysis

Statistical analyses were performed using GraphPad Prism 3.0 (GraphPad Software).

## Acknowledgements

We would like to thank Dr. Jeremy Stark (COH, Duarte, CA, USA) for providing us the U2OS SA-GFP cells (containing the SSA reporter), Dr. Maria Jasin (MSKCC, MYC, NY, USA) for the ATPm-RAD51s expression vectors and Dr. Gaelle Legube (Toulouse, France) for sharing the DIvA systems and technology. This work was supported by grants from the Ligue Nationale contre le cancer “Equipe labellisée 2017”, ANR (Agence Nationale de la Recherche, ANR-14-CE10-0010-02, ANR-16-CE12-0011-02, and ANR-16-CE18-0012-02), AFM-Téléthon and INCa (Institut National du Cancer, 2013-1-PLBIO-14).

## Author contributions

Conceptualization: JG-B, BSL; Funding acquisition: BSL; Investigation: JG-B, AM, CC, LCC, SR, YC, PD; Methodology: JG-B, ELC, JYM, PD, BSL; Project administration: BSL; Resources: YC, ELC, JYM, BSL; Supervision: JG-B, YC, ELC, JYM, PD, BSL; Validation: Protein purification; LSS, JYM; Writing original draft: JG-B, BSL; Writing – review & editing: JG-B, YC, JYM, ELC, PD, BSL.

## Supplementary Material

**Supplementary data S1:**
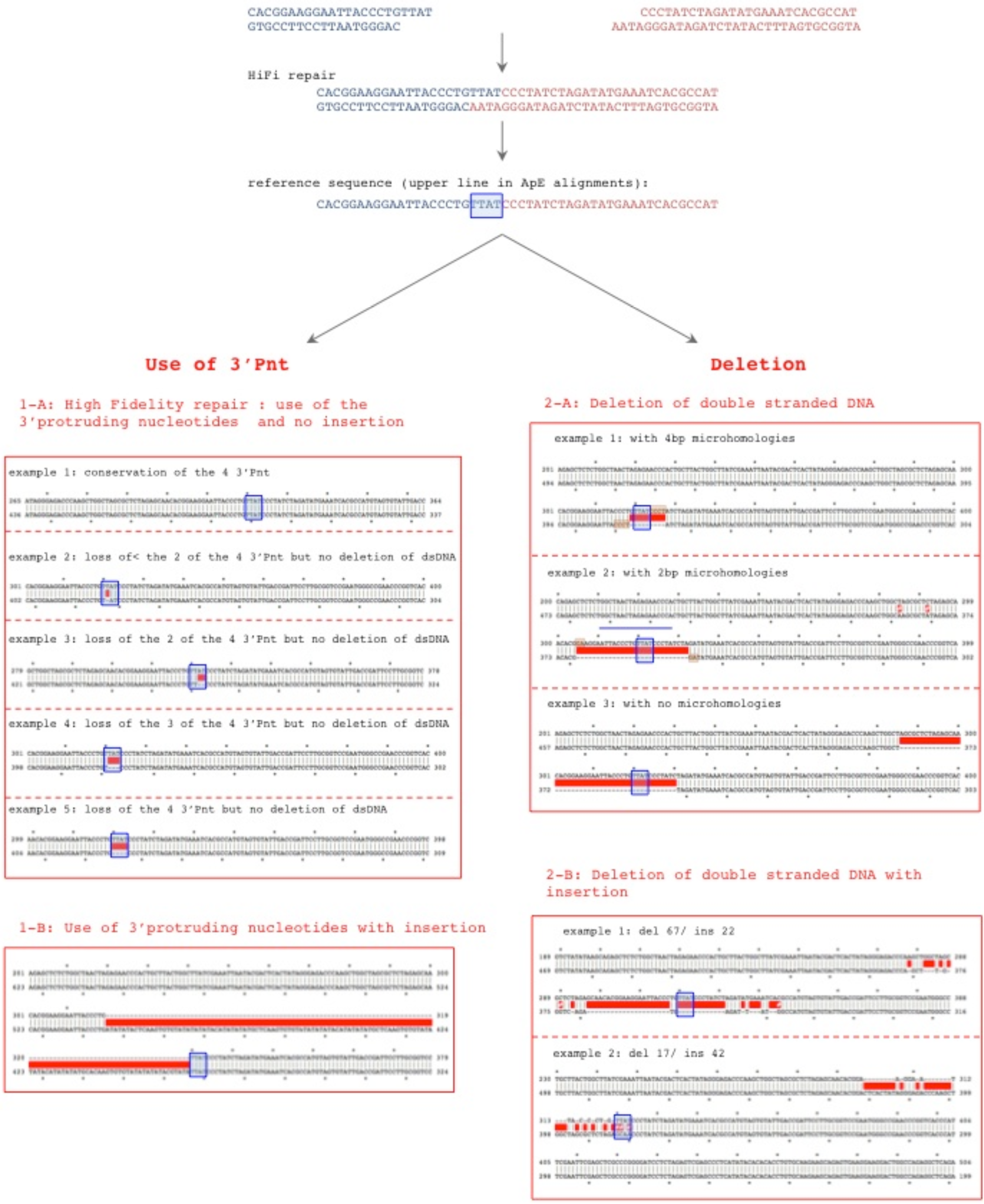
Alignments of repair scars with the expected sequence. Alignments are performed with the Ape software. Left panel : Use of 3’Protruding nucleotides (3’Pnt), no resection of the double stranded DNA. **1-A**: Use of the 3’ Pnt; **1-B** : the 3’PNT are present in the repair scar but an additional ectopic sequence is inserted. Right panel : repair scars with resection of the double stranded DNA leading to a deletion at the scar. **2-A**: Simple deletion. Microhomologies can eventually be used at the junction (highlighted in pink) **2-B**: A deletion can be coupled to the insertion of a non aligned/ectopic sequence.

**Supplementary data S2: Sequencing data of repair scars on the CD4-3200bp reporter in cells transfected with the different siRNA** See Excel file

**Supplementary data S3:**
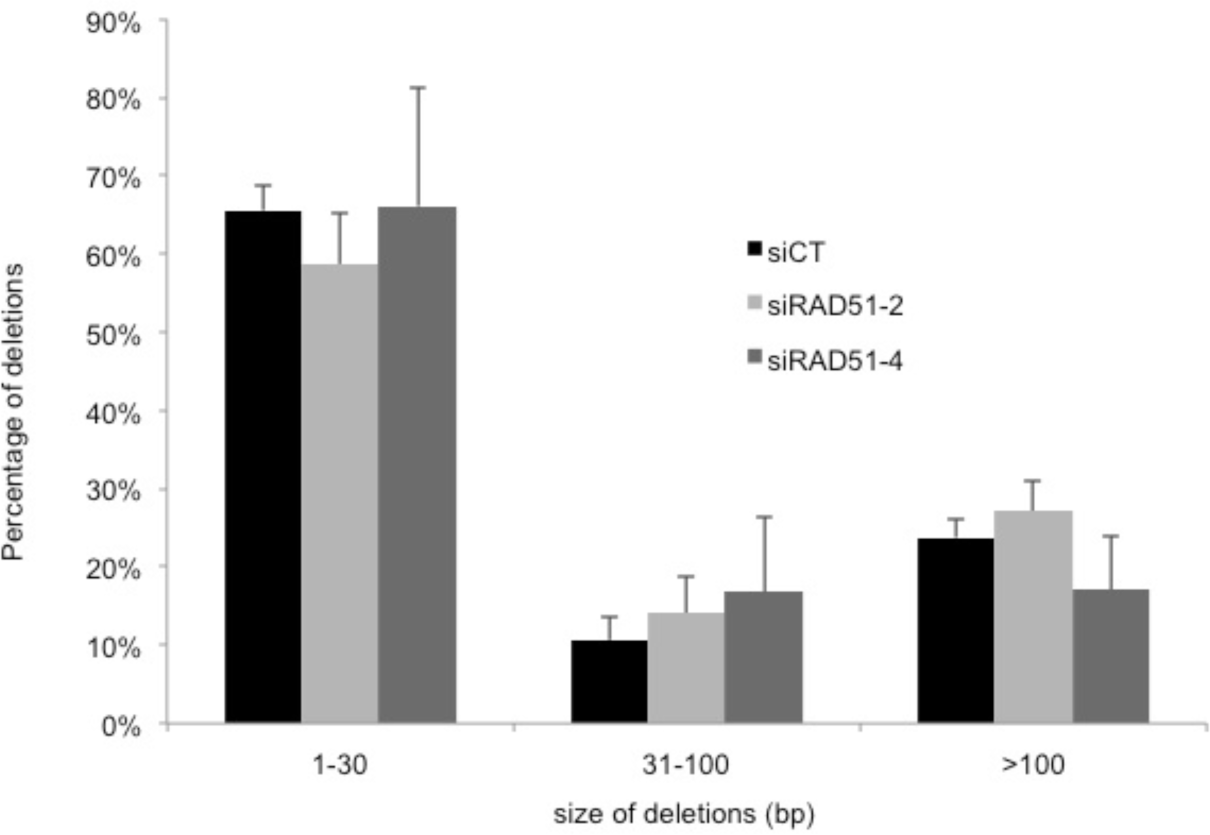
Distribution of the deletions size measured in cells bearing the CD4-3200bp reporter upon silencing of RAD51 with 2 different siRNAs. The values represent the average ± SEM of 2 to 5 independent experiments.

